# Floristic variation along landscape transformation gradients in successional tropical dry forests of the Colombian inter-Andean valleys

**DOI:** 10.64898/2026.08.02.741860

**Authors:** Ana Maria Reyes-B, Daniel Hernán Garcia Villalobos

## Abstract

This study analyzed variation in the floristic composition of successional tropical dry forests (Bs-T) along a gradient of landscape transformation and environmental factors in the Colombian inter-Andean valleys. Using 45 permanent plots distributed across the Magdalena and Cauca river basins, floristic patterns were evaluated through analyses of diversity (Hill numbers), similarity (NMDS), and relationships with environmental variables (RDA). The results showed a clear floristic differentiation between watersheds, whereas the succession and transformation gradients were not significant in explaining compositional variation. The main variables associated with floristic composition were total annual precipitation, mean annual temperature, and edaphic properties (pH, texture, base content, and available phosphorus), showing that environmental and geomorphological heterogeneity is the main determinant of floristic patterns in these ecosystems. Although some plots exhibit structural features typical of late successional stages, floristic composition reflects species assemblages characteristic of early and intermediate stages, suggesting that Bs-T forests require periods exceeding 50 years to recover the typical composition of mature forests. Taken together, these results confirm that floristic variation in the inter-Andean valleys is primarily related to multiscale environmental gradients rather than to the direct effects of transformation or succession, highlighting the need to incorporate these factors into management, restoration, and conservation strategies for tropical dry forest.

## Introduction

At the global level, more than 6.4 million hectares per year of tropical forests were degraded between 1990 and 2015, mainly as a result of anthropogenic activities such as the expansion of the agricultural frontier, mining, and infrastructure development (Keenan *et al.,* 2015). Part of these transformed areas are abandoned, and as a result of the natural regeneration process, successional forests develop in them, following diverse trajectories depending on site-specific factors such as the type of transformation and environmental variables, among others (Arroyo-Rodríguez *et al.,* 2017). As a consequence of this anthropogenic intervention, more than 50% of tropical forests are now successional (Chazdon *et al*., 2016), which makes the study of these forests increasingly relevant Chazdon, 2017; Hurtado-M *et al.,* 2021; Pérez-Cárdenas *et al*., 2021).

Understanding the floristic composition of secondary forests under different transformation scenarios is fundamental to understanding their variation along successional trajectories (Guariguata *et al.,* 2001; Lebrija-Trejos *et al*., 2010). This requires a dynamic assessment of diversity, considering that it is not a static factor but rather changes across scales (temporal, geographic, and biotic) (Mori *et al*., 2017). Changes in floristic composition can move in two directions: toward homogeneity, explained through taxonomic simplification, in which the presence of generalist species adapted to the limiting conditions generated by disturbances increases, denoting a loss of the forest’s capacity to recover its original state (Thier *et al.,* 2016; Hurtado-M *et al.,* 2021; Yang *et al*., 2021). In contrast, there is floristic heterogeneity, which reflects the responsiveness of vegetation to disturbance, conditioned by factors including low connectivity between patches, as well as the large distances between mature forests and successional remnants, leading to high floristic differentiation of emerging vegetation among fragments (Franklin *et al.,* 2015; González-M *et al.,* 2018; Arroyo-Rodríguez *et al*., 2023).

Assessing floristic variation in the Tropical Dry Forest (bs-T) in Colombia is relevant because it is a vegetation formation strongly impacted by human activities such as livestock farming, agriculture, and, more recently, infrastructure development, which act as transforming agents of these forests (González-M *et al.,* 2018). About 78% of the bs-T remnants in the country correspond to successional forests (García & González-M, 2019), dominated by early and intermediate successional stages (González-M *et al.,* 2018). At the regional level, marked floristic heterogeneity has been documented, determined by the differentiation in the country’s geomorphological and environmental characteristics, as well as by the different disturbances, which vary in type, intensity, and frequency depending on the region and the scale of analysis (Castellanos-Castro & Newton, 2015; Dryflor *et al*., 2016; González-M *et al*., 2018).

The dry forests of the inter-Andean valleys associated with the Cauca and Magdalena river basins are characterized by their location in lowlands, where features such as generally basic, nutrient-rich soils and a climate with low rainfall provide the best conditions for agricultural and livestock practices (Espinal & Montenegro, 1977). Consequently, these territories lie within an agricultural-livestock matrix, with irregular forest patches at different successional stages (Arcila Cardona *et al.,* 2012; Pizano *et al*., 2014; Rodríguez & López, 2014; González-M *et al.,* 2018). Due to the complexity of the transformed landscape, it is important to consider both land-use history and climatic characteristics at the regional and local scale (Arroyo *et al.,* 2015). These factors can determine the number and size of forest remnants in the region and, consequently, the species pool involved in the responsiveness of successional vegetation to different disturbances (Ewers *et al.,* 2013; Arroyo *et al.,* 2015).

This research aims to analyze variation in the floristic composition of successional dry forests along gradients of landscape transformation and environmental factors in the inter-Andean valleys of Colombia. Specifically, it addresses the following questions: (1) How does floristic composition differ across spatial scales and levels of landscape transformation? and (2) To what extent are climatic and edaphic conditions related to floristic variation in successional dry forests? We hypothesize that floristic composition exhibits marked heterogeneity at all scales of analysis, resulting from the influence of local environmental and edaphic conditions. However, in more transformed landscapes and early successional stages, the predominance of generalist species shared among windows and watersheds is expected, leading to a partial reduction in floristic differentiation. Understanding these patterns will contribute to the knowledge of the ecological processes that determine vegetation organization in tropical dry forests and help identify the factors that may favor or limit their recovery under land-use change scenarios.

## Methodology

### Study area

This research was conducted in the geomorphological unit of the inter-Andean valleys, defined by the Cauca and Magdalena river basins. The 45 permanent tropical dry forest (Bs-T) monitoring plots considered in this study are distributed across these basins, located in four departments: Huila and Tolima, belonging to the upper Magdalena river basin, and Valle del Cauca and Antioquia, associated with the upper and lower Cauca river basin, respectively (Figure 1).

**Figure 1.**
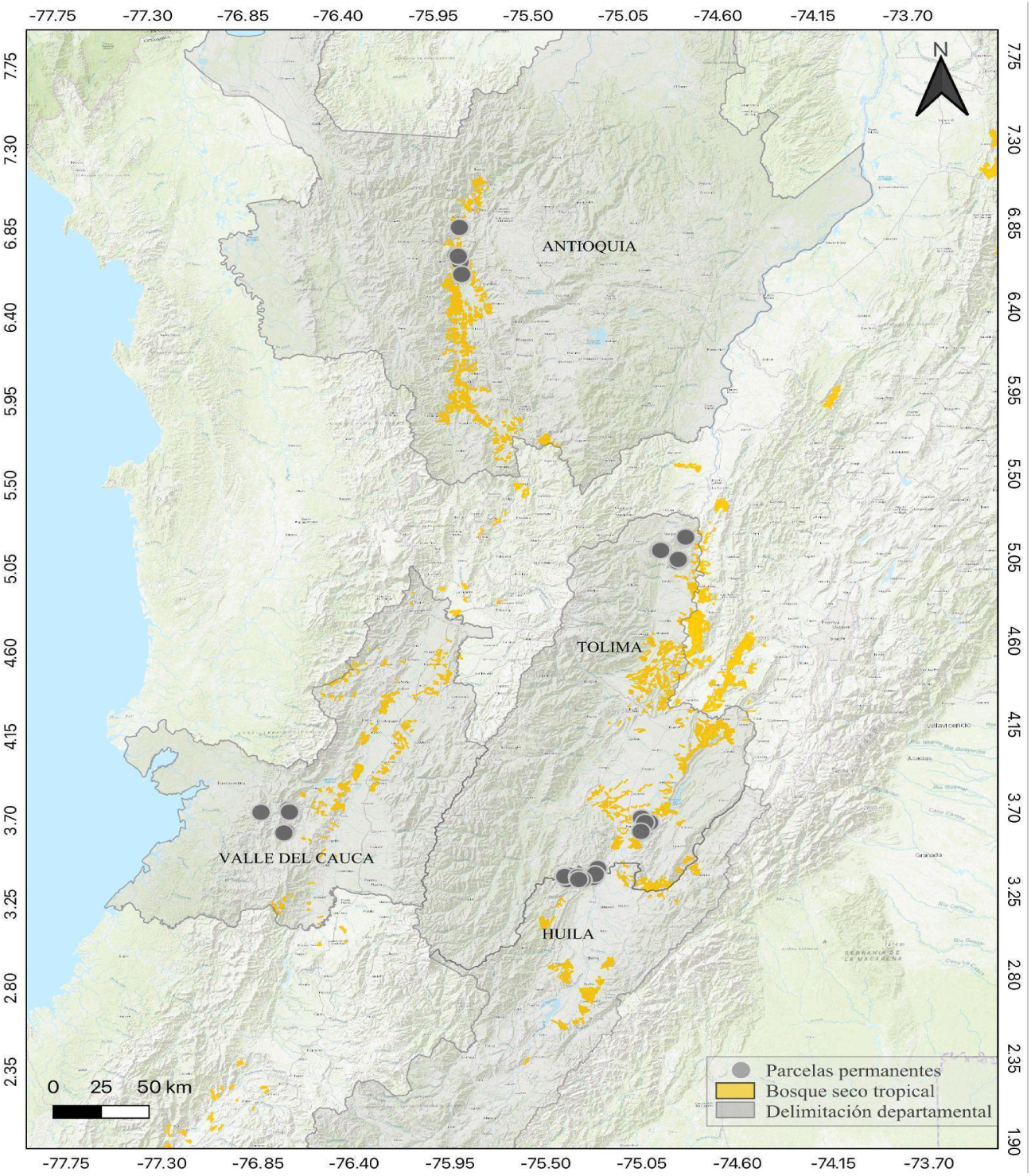
Location of the permanent plots in the 4 departments: Antioquia, Valle del Cauca, Huila, and Tolima. The distribution of Bs-T in the study area is shown in yellow.

The inter-Andean valleys have an average annual precipitation of 1535 to 2185 mm; however, the greatest differentiation between watersheds occurs toward the middle zone of the mountain ranges: in the foothills of the Western and Central Cordilleras, precipitation reaches up to 4500 mm, whereas in the valley of the Eastern Cordillera, from the department of Huila to the Cundiboyacense highlands, precipitation ranges between 500 and 1500 mm (Arango *et al.,* 2013a). In general, the inter-Andean valleys have two dry periods (December-March and June-September) and two wet periods (March-May and October-November), and a mean annual temperature of 26°C with diurnal variations of more than 10°C (Restrepo, 2005).

The Cauca river basin, bounded by the Central and Western Cordilleras, is characterized by an alluvial plain with low slopes (<5°) in the upper zone and a narrow valley toward the middle and lower, northward-flowing section with steep slopes (>35°) (Restrepo, 2005). On the other hand, the upper Magdalena river basin is described as a depression extending from Pitalito (Huila) to Honda (Tolima). It is a narrow valley with steep slopes (>35°) that generate an extensive drainage network, which intensifies the erosive process on the slopes of the Central and Eastern Cordilleras (Restrepo, 2005).

### Plot establishment

This research was based on secondary information derived from censuses conducted in the permanent monitoring plots, established between 2016 and 2017 within the framework of projects carried out by UNDP and IDB, under the leadership of the Humboldt Institute. The plots are distributed across two regions: the geographic valley of the upper Magdalena river basin, in the municipalities of Honda, Falan, Armero Guayabal, Natagaima, and Aipe; and the geographic valley of the Cauca river basin, in the municipalities of Buriticá, Liborina, Sabanalarga, Santafé de Antioquia and Dagua (Table 1).

**Table 1.** Distribution of the permanent plots across the inter-Andean valley watersheds, organized by department and municipality, indicating the landscape transformation level and the early (Tem), intermediate (Int), and late (Tar) successional stages evaluated.

| **Watershed** | **Department** | **Municipality** | **Transformation** | **Succession** |
| --- | --- | --- | --- | --- |
| **Cauca** | Antioquia | Buritaca, Laborina | High | Tem, Int, Tar |
|  |  | Santafé de Antioquia | Medium |  |
|  |  | Sabanalarga | Low |  |
|  | Valle del Cauca | Dagua | High, medium, and low |  |
| **Magdalena** | Tolima norte | Falan | High | Tem, Int, Tar |
|  |  | Honda | Medium |  |
|  |  | Armero Guayabal | Low |  |
|  | Tolima sur | Natagaima | High, medium, and low |  |
|  | Huila | Aipe | High, medium, and low |  |

Plot placement was based on two main criteria: the landscape transformation gradient (high, medium, and low) and the successional gradient (early, intermediate, and late) (Salgado-Negret et al., 2017). The selection of the study areas considered relief, hydrology, percentage of vegetation cover, and land-use change in transformed landscapes (Salgado-Negret et al., 2017). Each area corresponds to a landscape window, defined in this study as a spatial unit of analysis that groups a set of plots systematically represented across landscape transformation gradients and successional stages. Five landscape windows were established, three in the Magdalena basin and two in the Cauca basin. Each window is made up of nine plots, organized into three transformation levels (high, medium, and low), within which a successional trajectory composed of three stages (early, intermediate, and late) is established (Table 1).

In each window, 3 sub-areas were established to define the transformation levels based on the area of remaining forest patches, classifying them as follows: high transformation, corresponding to a matrix with a low proportion of remaining patches, high fragmentation, and low recovery; medium transformation, determined by the midpoint of patch proportion, size, and connectivity; and low transformation, where the matrix consists of a higher proportion of size and connectivity of forest patches that remain or are recovering (Salgado-Negret et al., 2017).

To determine the three successional stages, the time of forest persistence, recovery, and disappearance over a 23-year period (1990-2013) was considered; based on this criterion, each patch was classified into the successional stage of late, intermediate, or early cover (mature forest, secondary forest, and scrubland) (Salgado-Negret *et al.,* 2017). Late-stage cover is defined as the area made up of a plant community with predominantly tree elements, forming a more or less continuous canopy, with cover representing more than 70% of the total area (IDEAM, 2008). These are little or non-intervened vegetation formations, whose original structure and functional characteristics have not been altered. Intermediate cover is defined as an open plant formation with typically tree elements whose canopy is discontinuous and represents between 30% and 70% of the total area. It is characterized by low recent-use intervention; its structure and composition may derive from disturbed cover that regenerated naturally over a period exceeding 23 years, with a persistence process of more than 4 years. Early cover, defined as secondary vegetation cover, results from the natural successional process following a disturbance event or destruction of the original vegetation. Its regeneration process spans a period of less than 4 years (Salgado-Negret *et al.,* 2017).

### Sampling areas

Plots of 0.1 ha (20 m x 50 m with 10 m x 10 m quadrants) were installed for each successional stage at each transformation level. Dasometric height data were recorded for all trees with a diameter greater than 2.5 cm at 130 cm height, along with the taxonomic determination of the species present based on botanical specimens, reviewed by specialists using herbaria and following the APG IV taxonomic classification system (APG IV, 2016)

### Data analysis

The floristic richness of each plot along the successional and landscape transformation gradient was characterized by fitting the observed richness to sample size, represented by the Hill diversity numbers q0, q1, q2 (q0=richness, q1=Shannon exponential index, and q2=inverse Simpson index) (Hill, 1973). Hill numbers provide a simple, easy-to-understand measure of species diversity in a given area, since they allow the information on species composition to be summarized in a single number (Ohlmann et al., 2019), which facilitates comparison among different sites and among succession and landscape transformation factors.

Floristic composition was analyzed using non-metric multidimensional scaling (NMDS), based on the Bray-Curtis dissimilarity index. This non-parametric method allows floristic dissimilarity patterns to be represented in a low-dimensional space, preserving the ecological relationships among communities (Legendre & Legendre, 2012). NMDS has been recommended in tropical vegetation studies for its ability to handle data with high variability and a large number of zeros (Clarke, 1993; McCune & Grace, 2002), and has proven to be an effective tool for identifying floristic groupings associated with environmental and disturbance gradients (Balvanera & Aguirre, 2006; Avella et al., 2019). The significance of floristic differences between successional and transformation levels was verified through a PERMANOVA using 999 permutations. This approach allows multivariate hypotheses to be evaluated without requiring assumptions of normality and is widely applied in community ecology studies (Anderson, 2001; De Souza *et al*., 2017). These were analyzed using the vegan package (Oksanen et al, 2022) in RStudio software (2021).

To characterize the vegetation structure across the different successional states evaluated, two complementary approaches were used: analysis of the Importance Value Index (IVI) and determination of vertical structure through the stratification criterion. The Importance Value Index (IVI) was calculated for each recorded species, integrating the parameters of relative density, relative frequency, and relative dominance (expressed through basal area), following the classic methodology of Curtis & McIntosh (1951). This index allows the identification of species with the greatest ecological weight within the plant community, reflecting their structural contribution and their role in canopy organization at each successional stage.

For the vertical structure analysis, the stratification criterion proposed by Rangel-Ch & Lozano (1986) was adopted, which defines vegetation strata based on height ranges established for tropical ecosystems. Based on this criterion, each recorded individual was assigned to its corresponding stratum: shrub (1.5 - 4.9 m); small trees (5 - 11.9 m); lower tree layer (12 - 25 m); and upper tree layer (> 25 m). The relative frequencies of each stratum were calculated based on the number of individuals present. This analysis made it possible to describe the vertical organization of the plant community and establish comparisons between the different successional states, revealing changes in structural complexity associated with the progress of ecological succession.

### Climatic, edaphic, and landscape variables

Climatic variables were obtained by processing WorldClim version 2.1 data (https://www.worldclim.org/) values corresponding to a historical monthly average over the 1970-2000 period. These data were obtained using the geodata package (Hijmans *et al.,* 2024) in the R programming language (R Core Team, 2021). This global repository provides climate layers interpolated from meteorological data from ground stations, offering a spatial resolution of up to 30 arc-seconds (∼1 km^2^), suitable for ecological analyses at the regional and landscape scale (Fick & Hijmans, 2017). Values corresponding to the location of each sampling plot were extracted from these layers using geographic coordinates. The variables considered were: mean annual temperature (TMA), mean temperature of the warmest month (TMMC), total annual precipitation (PTA), and precipitation of the driest month (PMS). These variables represent fundamental thermal and hydric gradients for understanding the distribution and floristic assembly of tropical dry forests (Borchert, 1994; Powers *et al.,* 2018).

The edaphic variables used in this study were pH, total organic carbon (totalCO, %), nitrogen (N, %), potassium (K), magnesium (Mg), sodium (Na), available phosphorus (Pdis), and the textural fractions of sand, silt, and clay (%). These data come from the research of González-M *et al*. (2019), who obtained the edaphic information from 10 soil samples randomly taken at each plot and analyzed following standardized protocols. The selected edaphic variables reflect nutrient availability, water retention, and soil chemical conditions that influence resource uptake, physiology, and plant adaptive strategies to water stress and fertility in tropical dry forests (Van Der Putten *et al.,* 2013; van der Sande *et al.,* 2023).

Finally, the landscape cover and transformation categories used in this study were taken from Salinas (2025), who classified areas based on satellite images (2014–2017) using supervised interpretation within a 500 m radius around each plot. In that work, cover types were classified as forest (Forest), secondary vegetation (SecVeg), and other human uses (UCL), and additional metrics of shape and terrain roughness were calculated in ArcGIS 10.8 (ESRI, 2020). The author shared the resulting data, which were used directly in the analyses of this study.

To identify the environmental variables associated with floristic composition, a Redundancy Analysis (RDA), which considers the edaphic, climatic, and landscape attributes as predictors. RDA allows linear relationships to be established between species abundance and environmental gradients (Ter Braak, 1986; Legendre & Legendre, 2012), a tool widely used to interpret floristic patterns in tropical ecosystems (Dexter *et al*., 2015; Aguirre-Gutiérrez *et al.,* 2020).

## Results

### Floristic richness and diversity

The regional floristic composition comprises 61 families, 190 genera, and 367 species. The most abundant families were Myrtaceae, with 2293 recorded individuals, followed by Fabaceae (1054 individuals) and Malvaceae (715 individuals). At the watershed level, in the Cauca valley, Myrtaceae with 987 individuals stands out, followed by Phyllanthaceae (530 individuals). Meanwhile, in the Magdalena watershed stand out Myrtaceae (1306 individuals) and Fabaceae (738 individuals).

Based on the diversity analysis using Hill numbers, it is evident that the Magdalena watershed showed higher richness (*q_0_* = 276) compared to the Cauca watershed (*q_0_* = 134). Likewise, the distribution of abundances was more even in the Magdalena (q₁ = 88; *q_2_* = 43), whereas the Cauca showed greater dominance by a few species (q₁ = 37; *q_2_* = 19). However, the non-parametric Kruskal-Wallis analysis indicated that species richness (*q_0_)* did not differ significantly between watersheds (p = 0.317).

Analysis of the successional gradient by watershed reveals differentiated patterns in the dynamics of species richness and evenness. In the Magdalena watershed, species richness increases progressively along succession, reaching its maximum in the late stages (q₁ = 158). However, evenness was higher in the intermediate stage, suggesting that at this phase the community reaches a more balanced structure, characterized by a more homogeneous distribution of dominance among species. In contrast, in the Cauca watershed, the intermediate and late successional stages showed similar richness (*q*₁ *=* 81 and 82, respectively), while the early stage showed considerably lower values (*q*₁ *=* 60). Even so, the results of the Kruskal–Wallis test indicated that differences in richness between successional stages were not statistically significant in either watershed (Magdalena: *p =* 0,4; Cauca: *p =* 0,09.

Regarding the landscape transformation gradient, contrasting responses were observed between watersheds. In the Cauca, richness increased as the transformation level decreased; conversely, in the Magdalena an opposite trend was evident: sites with a higher level of transformation showed greater species richness.

Non-metric multidimensional scaling (NMDS) analysis made it possible to explore patterns of variation in floristic composition among plots. The two-dimensional model showed a *stress* value of 0.20, indicating an acceptable representation of floristic similarities in the reduced space. The results of the fit of the factors explored through the analysis showed a significant relationship between watershed and floristic composition (*R^2^* = 0,07, *p* = 0,001), reflecting a slight floristic differentiation between both regions. In contrast, the succession (*R^2^* = 0,04, *p* = 0.737*) and landscape transformation (*R^2^* = 0.05, *p* = 0.133*) factors showed no statistically significant effects. Graphically, plots associated with the Cauca watershed tended to group toward the upper right quadrant of the NMDS plane, while those from the Magdalena were concentrated toward the central and lower left sector, which is consistent with the pattern of differentiation detected statistically (Figure 1). No defined groupings were observed according to successional stages (early, intermediate, and late).

At the watershed scale, in addition to the succession and transformation factors, the landscape window factor associated with each watershed was incorporated. Similar to the regional analysis, a significant dissimilarity pattern was detected (R^2^ = 0.18, p = 0.001; R^2^ = 0.13, p = 0.017) associated with the landscape windows in each watershed (Figure 2a and 2c). In contrast, the landscape transformation factor showed no significant effect on the configuration of plant communities (R^2^ = 0.08, p = 0.335; R^2^ = 0.12, p = 0.414).

**Figure 2.**
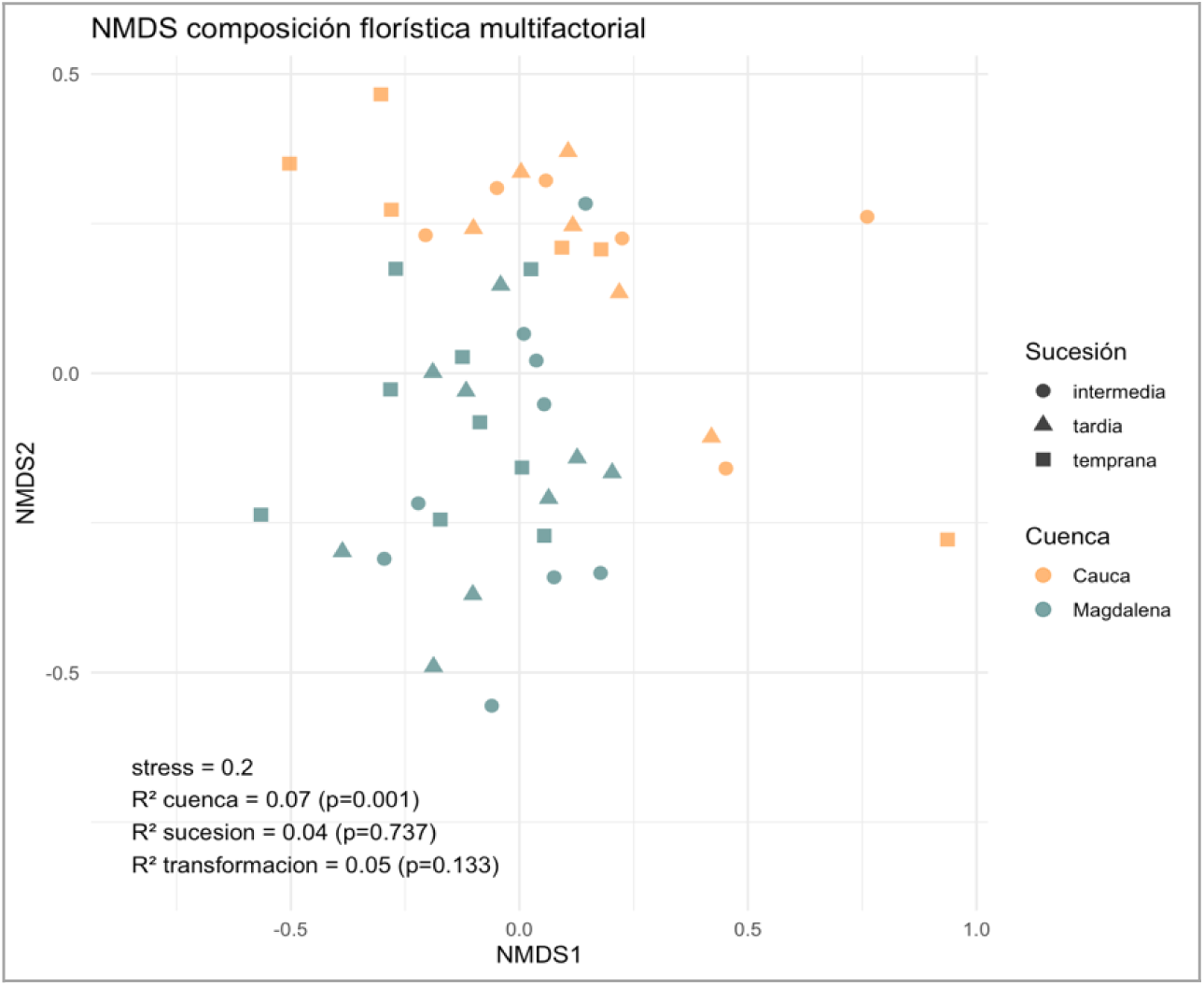
Ordination space of the floristic composition of 45 plots located in two watersheds of the inter-Andean valleys (Cauca and Magdalena river basins) based on the Bray-Curtis similarity index, as a function of watershed, successional gradient, and transformation level.

Spatially, in the Magdalena watershed a grouping is observed in the upper right quadrant of the plots corresponding to the Tolima S ur and Huila windows, suggesting a greater similarity in species composition between these windows. In contrast, the plots of the Tolima Norte window are concentrated toward the upper left quadrant and the center of the plot, evidencing the differentiated floristic composition. For the Cauca watershed, the plots associated with each transformation window group according to this scale, where Antioquia groups along the positive axis of NMDS 1, while plots from Valle del Cauca group to the left of the same axis. In contrast, the two-dimensional model based on the transformation level shows no defined grouping patterns, and significance values are high, suggesting that local spatial variation contributes more strongly to the observed floristic heterogeneity (Figure 2 b and d).

**Figure 3.**
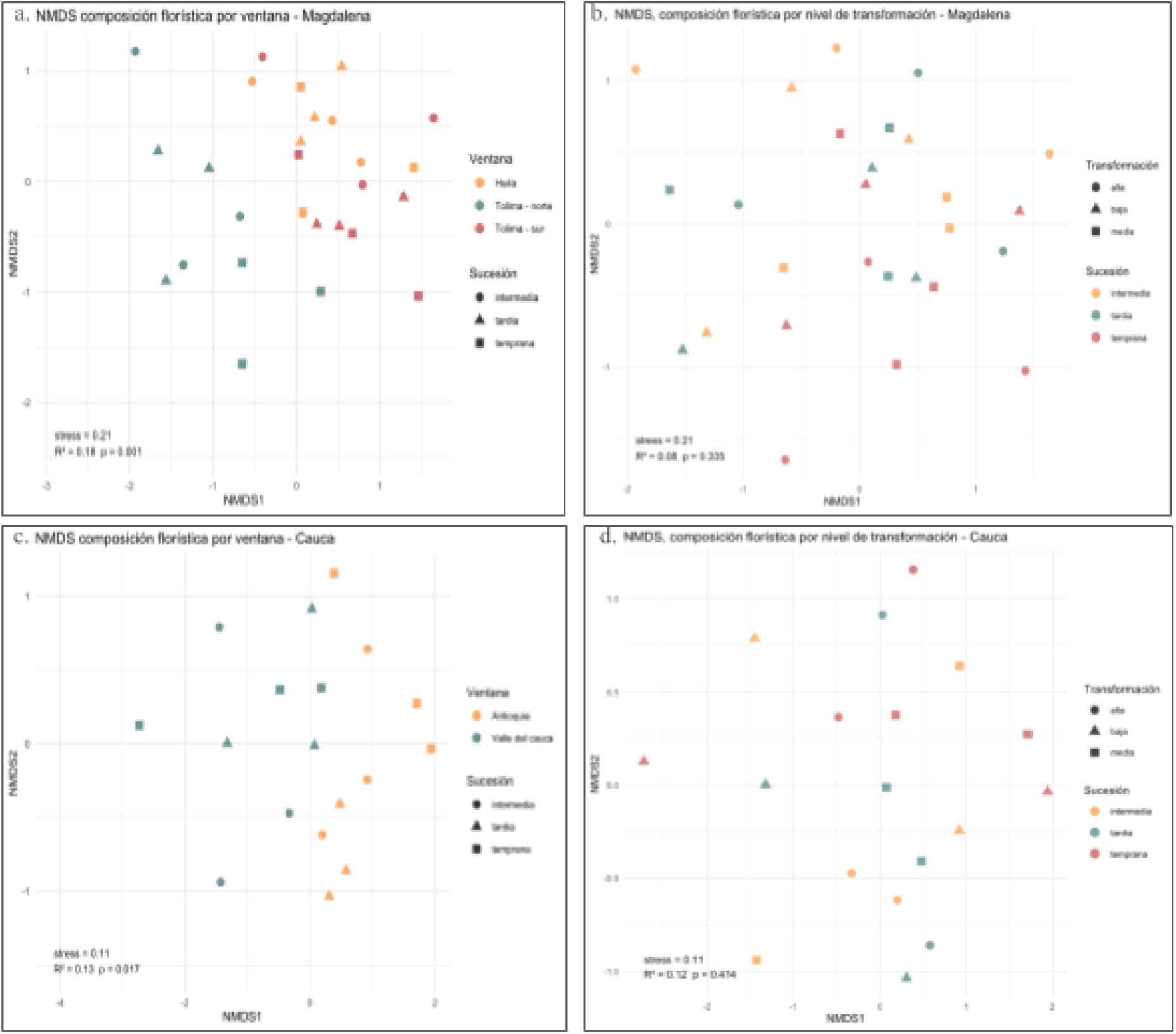
NMDS analysis of the floristic composition in the Magdalena and Cauca river basins. Panels a and c show the variation in floristic composition among landscape windows in each basin. In the Magdalena (a, b) and Cauca (c, d) basins. Panels a and c show significant segregation among landscape patches: in the Magdalena (a) between Huila, northern Tolima, and southern Tolima (stress = 0.11, R^2^ = 0.18, p = 0.001), and in the Cauca (c) between Antioquia and Valle del Cauca (stress = 0.21, R^2^ = 0.13, p = 0.017). Panels b and d show the effect of landscape transformation and successional stage as factors, without revealing any significant segregation pattern in any of the watersheds (Magdalena: R^2^ = 0.08, p = 0.335; Cauca: R^2^ = 0.12, p = 0.414). The symbols indicate successional stage, and the colors represent landscape windows (a, c) or levels of transformation (b, d).

Analysis of the most abundant species revealed differentiated dominance patterns between watersheds as a function of landscape transformation level and successional stage (Figure 5). In the Cauca watershed, the most abundant species varied according to the transformation gradient: in landscapes with high transformation, *Guettarda malacophylla y Guazuma ulmifolia* had the highest abundances, concentrated mainly in early and intermediate successional stages. In contrast, in landscapes with low transformation, *Phyllanthus botryanthus y Myrcia fallax* dominated the intermediate and late stages, whereas at medium transformation levels, *Phyllanthus botryanthus, Murraya paniculata* y *Malpighia glabra* showed a greater presence, distributed across the different successional stages.

**Figure 4.**
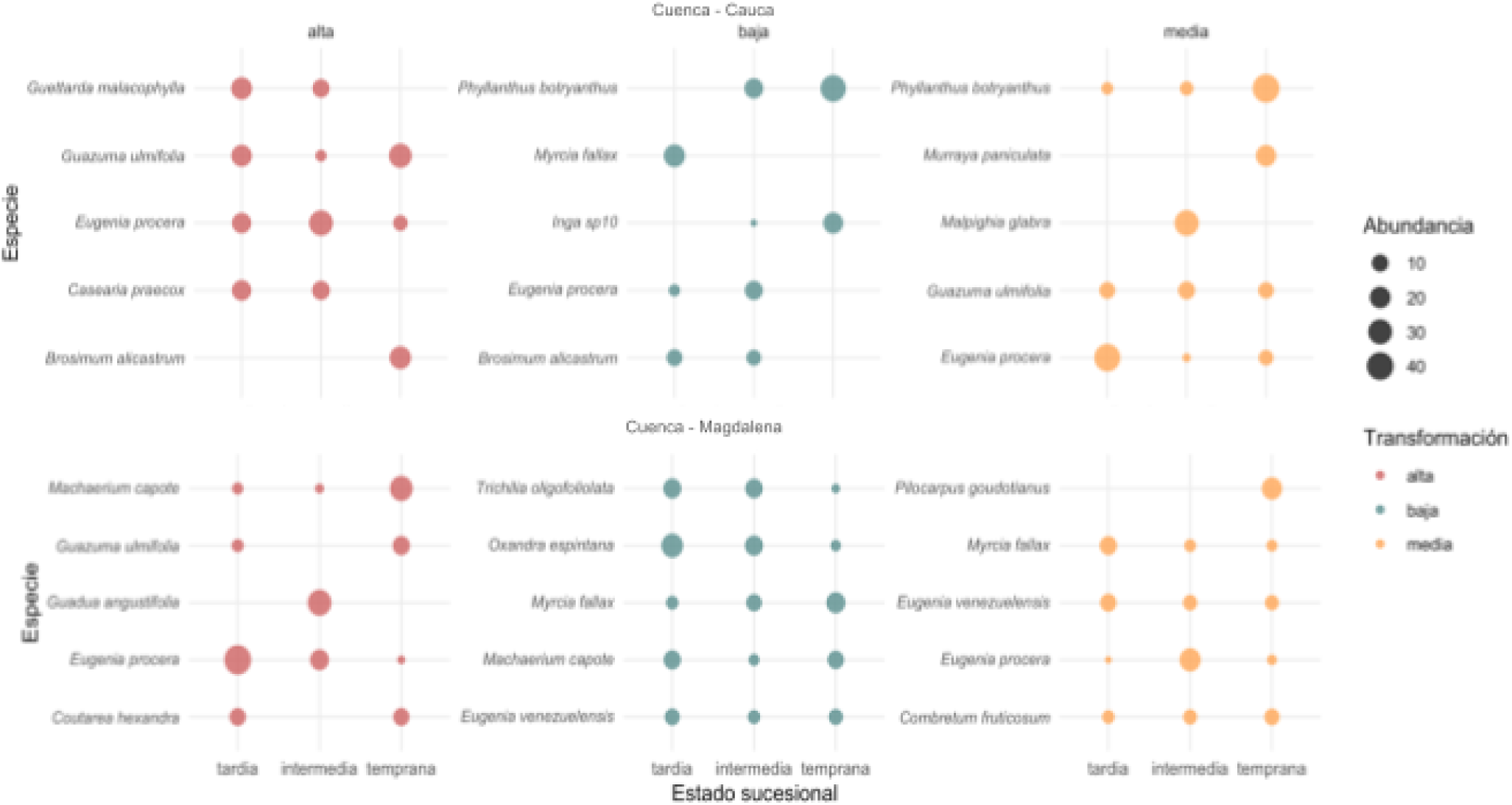
Abundance of the five most dominant species according to landscape transformation level and successional stage in the Cauca and Magdalena watersheds. Circle size indicates relative abundance and color represents transformation level.

In the Magdalena watershed, the dominance pattern was also influenced by landscape transformation. In landscapes with high transformation, *Machaerium capote y Guazuma ulmifolia* were the most abundant species, dominating especially in early stages and, to a lesser extent, in intermediate stages. In landscapes with low transformation, *Trichilia oligofoliolata, Oxandra espintana* y *Myrcia fallax* showed consistent abundances across the three successional stages, evidencing a more homogeneous distribution. For their part, in landscapes with medium transformation, *Eugenia venezuelensis, Myrcia fallax* y *Pilocarpus goudotianus* were the dominant species, with greater abundance in intermediate and early stages.

A pattern is evident in the behavior of *Eugenia procera*, which consistently appeared in multiple combinations of transformation and succession in both watersheds. This species showed considerable abundances both in highly transformed landscapes and in those with medium transformation, and was present in early, intermediate, and late stages. In contrast, species such as *Guettarda malacophylla* in the Cauca and *Machaerium capote* in the Magdalena showed a more specific association with highly transformed landscapes and early stages.

### Influence of environmental and landscape variables

The inter-Andean valleys have slightly acidic soils; the Cauca reaches values closer to neutrality (pH = 6.82) than the Magdalena (pH = 6.57). C and N contents are similar between watersheds, but the Magdalena records the highest values (0.25% versus 0.21%), while the Cauca stands out for a greater availability of bases (Mg, Na, and total bases), confirming greater edaphic fertility in this watershed (Table 2). Soil texture in both regions is clay loam, with a clay proportion close to 50%. The Cauca has a higher silt fraction, whereas in the Magdalena the soil is more clayey, implying differences in water and nutrient retention capacity. Phosphorus availability is high and similar between both watersheds, although with high variability.

**Table 2.** Mean values and standard deviation of the edaphic and climatic variables used for the redundancy analysis. (Data obtained and analyzed from the IDB and UNDP project)

| Variable | WATERSHED |  |
| --- | --- | --- |
|  | Cauca | Magdalena |
| pH | 6,82 ± 0,26 | 6,57 ± 0,20 |
| Total organic carbon (TotalCO) | 2,85 ± 0,86 % | 3,00 ± 0,81 % |
| Nitrogen (N_%) | $0,21 \pm 0,05 \%$ | $0,25 \pm 0,05 \%$ |
| Potassium (K) | $0,57 \pm 0,22 \text{ cmol.Kg}^{-1}$ | $0,50 \pm 0,10 \text{ cmol.Kg}^{-1}$ |
| Magnesium (Mg) | $7,40 \pm 3,67 \text{ cmol.Kg}^{-1}$ | $5,16 \pm 1,68 \text{ cmol.Kg}^{-1}$ |
| Sodium (Na) | $0,14 \pm 0,05 \text{ cmol.Kg}^{-1}$ | $0,07 \pm 0,03 \text{ cmol.Kg}^{-1}$ |
| Available phosphorus (avaiP) | $24,93 \pm 17,33 \text{ mg.Kg}^{-1}$ | $26,81 \pm 13,44 \text{ mg.Kg}^{-1}$ |
| Sand (sand_%) | $21,04 \pm 2,90 \%$ | $20,70 \pm 4,32 \%$ |
| Silt (Silt_%) | $28,78 \pm 2,70 \%$ | $27,56 \pm 4,51 \%$ |
| Clay (clay_%) | $49,47 \pm 6,14 \%$ | $50,68 \pm 9,43 \%$ |
| Bases (totalbases) | $32,72 \pm 2,98 \text{ cmol.Kg}^{-1}$ | $29,57 \pm 4,69 \text{ cmol.Kg}^{-1}$ |
| Altitude | $742.78 \pm 255.02 \text{ m a.s.l.}$ | $482.52 \pm 203.40 \text{ m a.s.l.}$ |
| Mean annual temperature (TMA) | $22,94 \pm 1,86 \text{ }^{\circ}\text{C}$ | $26,30 \pm 1,16 \text{ }^{\circ}\text{C}$ |
| Mean temperature of the driest month (TMMS) | $28,68 \pm 2,13 \text{ }^{\circ}\text{C}$ | $33,23 \pm 1,38 \text{ }^{\circ}\text{C}$ |
| Mean temperature of the driest quarter (TMTS) | $23,09 \pm 1,96 \text{ }^{\circ}\text{C}$ | $26,67 \pm 1,18 \text{ }^{\circ}\text{C}$ |
| Total annual precipitation (PTA) | $2117,17 \pm 546,18 \text{ mm}$ | $1906,19 \pm 269,32 \text{ mm}$ |
| Precipitation of the driest month (PMS) | $62,89 \pm 19,49 \text{ mm}$ | $59,74 \pm 21,31 \text{ mm}$ |

Climatic conditions are contrasting: the plots analyzed in the Cauca river valley watershed are located at a higher average altitude (743 m) than those of the Magdalena (483 m); this is reflected in lower mean temperatures (22.9°C versus 26.3°C, respectively). Consistently, the Cauca shows moderate maximum and minimum temperatures, as well as higher mean annual precipitation (2117 mm versus 1906 mm) (Table 2).

The redundancy analysis (RDA) showed a significant relationship between floristic composition and environmental variables (*R^2^ =* 0.64; *R^2^ad◻* = 0.22; *p =* 0.001). The first two axes jointly explain 16.1% of the total variance and reflect contrasting environmental gradients between the Cauca and Magdalena watersheds (Figure 5a). In the Cauca, forests were associated with soils with relatively high values of pH, Na, and clay; the most influential climatic variable is total annual precipitation (PTA); regarding landscape characteristics, the effect of the number of patches and forest area was found, evidencing the influence of landscape structure on floristic composition. In the Magdalena watershed, vegetation is divided according to soil composition, where plots with higher proportions of silt, sand, and total bases are found on the upper right axis, representing greater edaphic heterogeneity and possible water restrictions, while for plots located on the left axis, the climatic variables of precipitation (PMS) and temperature (TMA, TMTS, and TMMS), as well as the landscape variable (secondary vegetation cover – SecVeg), best explain the species composition in this watershed. These results show that the difference in floristic composition in these areas is mediated by regional environmental gradients.

**Figure 5.**
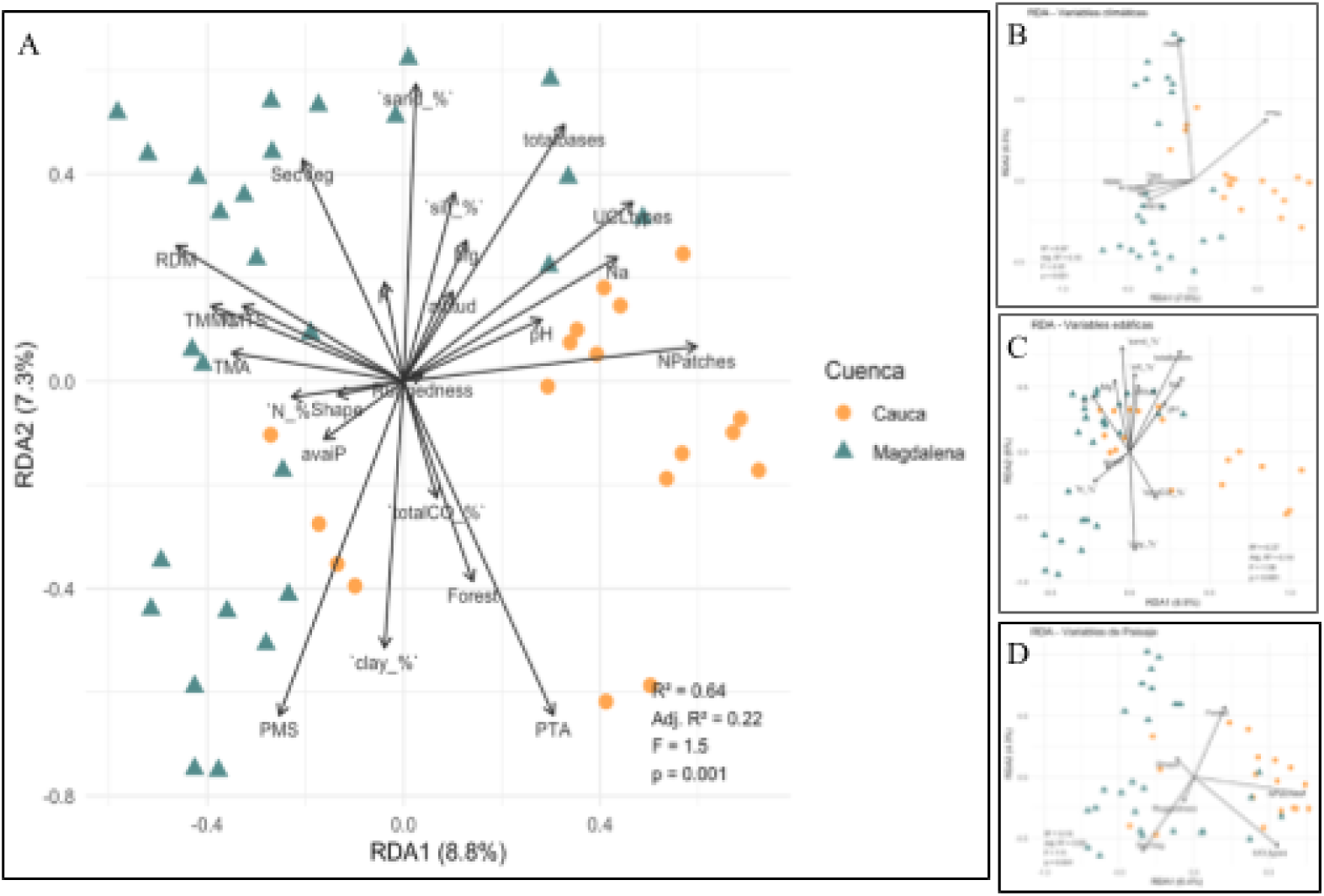
Redundancy Analysis (RDA) of floristic composition (A) as a function of climatic (B), edaphic (C), and landscape (D) variables for the Cauca and Magdalena watersheds.

At the watershed and landscape window scale, floristic variation patterns showed higher resolution in the response of composition to local environmental conditions. The RDA for the Cauca watershed (Figure 6a) showed a clear separation of plots according to window and associated environmental conditions. Climatic variables, such as total annual precipitation (PTA) and mean annual temperature (TMA), together with landscape variables (forest cover and number of vegetation patches), were mainly related to the plots of the Antioquia window, characterized by higher values of precipitation, temperature, forest cover, and landscape fragmentation. In contrast, the plots corresponding to the Valle del Cauca window were associated with altitude gradients, secondary vegetation cover, and edaphic variables, particularly texture, pH, and base content. The model explained 32% of the total variation (*Adjusted R^2^* = 0.32; *p =* 0.001), indicating a significant relationship between environmental variables and floristic composition.

The windows for the Magdalena watershed also differ in environmental and landscape characteristics particular to each site (Figure 6c). Tolima Norte shows an association with higher values of precipitation, temperature, and forest cover, whereas Huila responds to edaphic variables such as pH and total organic carbon (TotalCO%) and landscape variables such as secondary vegetation cover and altitude. The Tolima Sur transformation window is mainly associated with landscape factors such as the number of patches, evidencing a highly fragmented matrix. The model explained 15% of the total variation (*Adjusted R^2^* = 0.15; *p =* 0.001).

Given that differentiation was evident between the landscape windows of each watershed, environmental and landscape variables were evaluated within each of them in order to analyze the organization of plots as a function of environmental, edaphic, landscape, and successional-stage factors. The results showed no patterns associated with succession, while edaphic and environmental factors were the ones that best explained the floristic composition of plots at the landscape window scale ( *Adjusted R^2^* = 0.39; *p =* 0.004) (Figure 6b and 6d). The applied model (variable configuration) did not show a uniform fit for all windows; it was adapted according to the variables that best explained floristic composition in each case.

**Figure 6.**
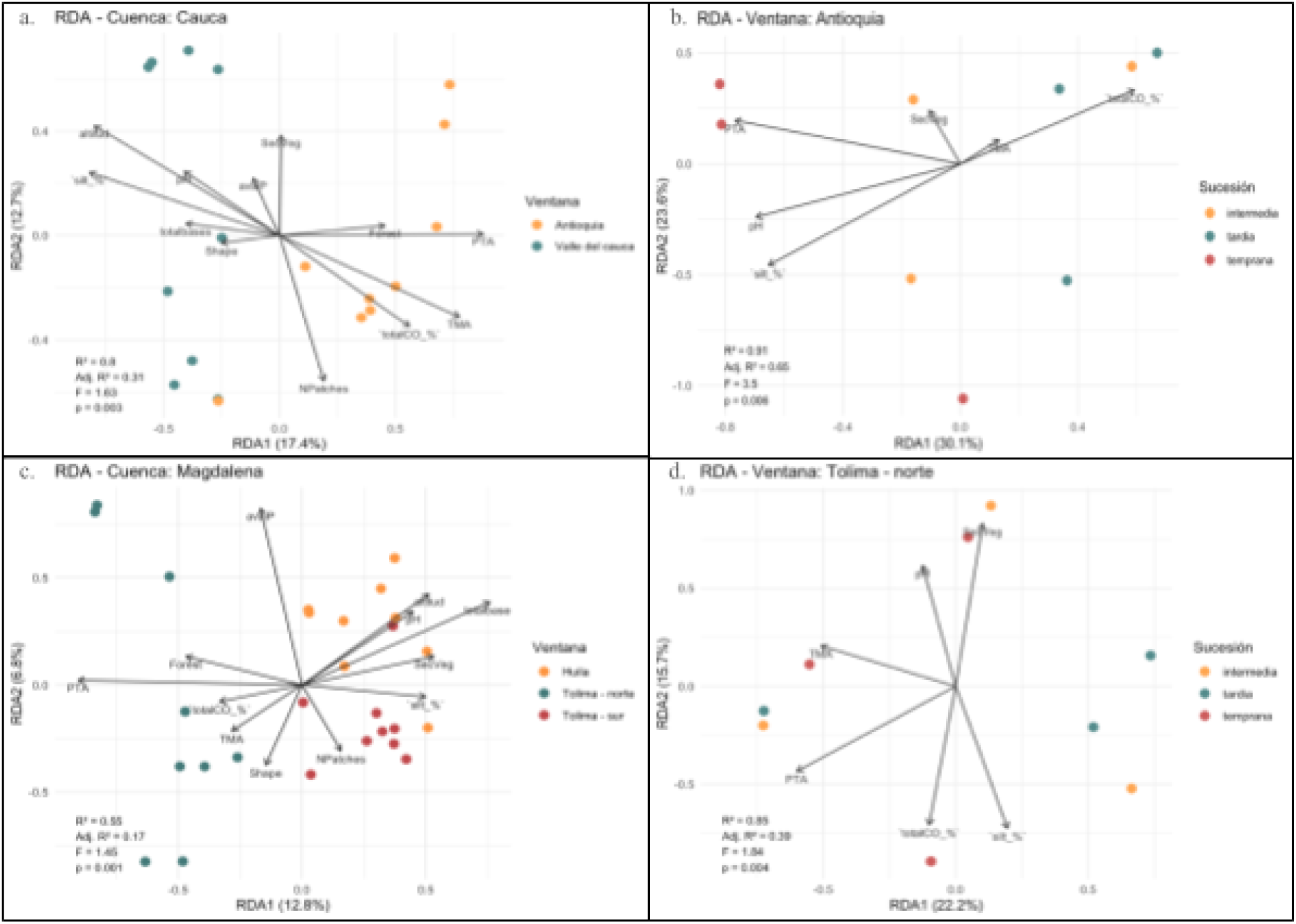
RDA of floristic composition according to environmental and landscape variables at the watershed (a, c) and window (b, d) scale in the inter-Andean valleys, as a function of climatic, edaphic, and landscape variables.

## Discussion

The tropical dry forest in the inter-Andean valleys shows high heterogeneity in its species composition; this variation is not directly related to the landscape transformation gradient or to successional stages. Instead, it appears to be related to ecological processes associated with local and regional environmental conditions. This suggests that floristic patterns do not respond linearly to transformation or succession gradients, but rather reflect the predominant influence of the environmental and biogeographic particularities of each landscape (Hernández *et al.,* 2011; Norden *et al.,* 2015).

Regional dissimilarity analysis using NMDS shows the segregation of plots according to watershed. The slight overlap observed between plots from both watersheds can be explained by the presence of widely distributed generalist species, while exclusive species reinforce the floristic identity of each watershed. Among the shared species, *Eugenia procera y Guazuma ulmifolia*, whose contrasting abundance patterns between regions and successional gradients reflect the heterogeneity of the ecological mechanisms that structure the vegetation (Figure 4).

Although the succession and transformation gradients showed no significant effects independently, their integrated analysis by watershed revealed differences in floristic composition between the Cauca and the Magdalena. *E. procera* showed high abundance across most successional stages and transformation levels, reflecting its broad tolerance to disturbance. However, in the Cauca its abundance decreased in less transformed landscapes with fewer patches, which could be related to its dependence on large-bodied dispersers (birds and flying mammals) that, given its berry-type fruit, tend to frequent open or edge areas (Gressler et al., 2006; Vargas, 2015). In contrast, *G. ulmifolia* was more abundant in the Cauca, especially in early stages and high and medium transformation levels, whereas in the Magdalena it predominated in early and intermediate stages under high transformation.

The dominance of these species could be related to their frequent incorporation into silvopastoral systems*. G. ulmifolia* is one of the tree species most widely used in these systems for its forage value: its leaves and fruits are palatable and digestible for livestock, which favors its conservation and dispersal in pastures (Giraldo, 1996). Dairy, beef, and dual-purpose cattle ranching is the predominant productive activity in the Colombian inter-Andean valleys, covering approximately 1,700,000 ha, with the Magdalena watershed being the region with the greatest livestock presence (Pulido et al., 1999). In this context, the persistence of *G. ulmifolia* in landscapes with high transformation and early successional stages would respond not only to its tolerance to disturbance, but also to active selection by producers, who retain them in pastures for the services they provide to the production system.

This behavior is consistent with their role as pioneer species in secondary succession processes, recognized in previous studies in tropical dry forests of the inter-Andean valleys (Torres-G et al., 2012; Vargas, 2015; Olascuaga-Vargas et al., 2016; Suárez & Vargas-R., 2019; Avella et al., 2019; Gerber et al., 2020). González et al. (2018) note that early and intermediate successional stages predominate in the tropical dry forests of these valleys, which explains why the most abundant species are characteristic of those stages. In this research, *E. procera y G. ulmifolia* obtained the highest Importance Value Indices (see appendices), were present in 24 of the 45 plots analyzed, and dominated the shrub (1.5–5 m) and small-tree (5–12 m) strata, characteristic of the early and intermediate successional stages in the inter-Andean valleys.

The coexistence of multiple successional stages within the same landscape increases beta diversity by generating structural and functional heterogeneity among patches, suggesting that different successional trajectories contribute to the maintenance of regional diversity even in highly transformed contexts (Avella et al., 2019; van Breugel et al., 2024). This pattern is consistent with the intermediate disturbance hypothesis, according to which richness and diversity increase in early and intermediate successional stages, where species from different stages coexist and the absence of equilibrium reduces the probability of competitive exclusion (Grime, 1973; Horn, 1975). However, landscape transformation and successional stage act as modulators rather than sole determinants of floristic composition: although they condition regeneration, recruitment, and dispersal (Arroyo-Rodríguez et al., 2017), their effect is mediated by local edaphic and climatic conditions.

In both watersheds, total annual precipitation (PTA) emerged as the most influential variable on floristic composition (Magdalena: F = 34.86, p = 0.001; Cauca: F = 35.98, p = 0.001), underscoring the dominant role of water availability in structuring tropical dry forests (Markesteijn et al., 2008; Allen et al., 2010; Castellanos-Castro & Newton, 2015; Aguirre-Gutiérrez et al., 2020). However, the variables that accompany and modulate this effect differed among windows within each watershed, evidencing differentiated ecological responses to environmental and landscape gradients: in the Magdalena watershed, the number of patches, base content, phosphorus availability, and mean annual temperature (TMA) were the most relevant explanatory variables (p = 0.028, 0.035, 0.37, 0.44, respectively), whereas in the Cauca, pH, TMA, and altitude stood out (p = 0.007, 0.011, 0.041, respectively). Consequently, the floristic structure and composition of the Magdalena appears more conditioned by water limitations and fragmentation processes, while that of the Cauca is more influenced by edaphic and altitudinal factors.

In contrast with what was reported by González-M *et al*. (2018), who identified the inter-Andean valleys as a relatively homogeneous unit in contrast to the other regions where bs-T is distributed in Colombia, the results of this research show marked floristic heterogeneity between the watersheds and windows analyzed. The low proportion of shared species suggests that the inter-Andean valleys do not form a uniform floristic set, but rather a mosaic of communities with particular compositions determined by local conditions and historical transformation processes. The apparent homogeneity detected in previous studies could be attributed to the influence of dominant species with wide geographic distribution, whose presence would tend to mask the underlying beta heterogeneity and the floristic particularities determined by the environmental and edaphic conditions of each watershed.

The climatic asymmetry can be explained, among other causes, by the uplift of the Central Cordillera, which acts as an orographic barrier to moisture flow from the Pacific, generating a dry shadow toward the Magdalena valley (Cuervo-Gómez et a., 2015; Salazar-Jaramillo et al., 2021). Consequently, these environmental contrasts, together with the geomorphological and biogeographic differences of each valley (Pennington et al., 2009; Dexter et al., 2015; González-M. et al., 2018; Maia et al., 2020; Hoorn et al., 2022), have favored ecological and floristic differentiation processes between both watersheds. The redundancy analyses and the magnitude of the environmental effects support the idea that spatial heterogeneity emerges as the dominant ecological mechanism, modulated by the interaction between edaphic gradients and local climatic conditions (Hernández-Stefanoni et al., 2011). Thus, edaphic variation—texture, fertility, and nutrient availability—explains much of the site-scale heterogeneity, while climatic gradients and biogeographic history sustain segregation at larger scales (Toledo *et al*., 2011; de Souza *et al.,* 2019; Maia et al., 2020).

These results allow two main conclusions to be drawn: first, no clear pattern is observed in floristic composition across successional stages or the different transformation levels, suggesting that in order to detect marked successional differences in these Bs-T, chronosequences including forests with longer recovery times than those considered in this study are likely required (Finegan, 1996); this is explained by the slow pace of successional processes under restrictive climatic conditions. Second, floristic differentiation cannot be attributed solely to succession or landscape transformation, but rather appears to be modulated by abiotic factors, especially water availability, temperature, and soil properties, whose influence was shown to vary hierarchically across scales. Consequently, conservation and restoration strategies must consider not only recovery time and landscape connectivity, but also the specific edaphic and climatic conditions of each watershed in order to promote effective functional and compositional recovery.

## Conclusions

The floristic composition of successional dry forests in the inter-Andean valleys is highly heterogeneous and does not follow a linear pattern with respect to succession or landscape transformation. The differentiation between watersheds reflects the predominant influence of local environmental and biogeographic factors, rather than the intensity of transformation or the successional stage.

At each scale, edaphic, climatic, and structural landscape gradients influence species composition differentially, reflecting the multiscale nature of the processes that structure inter-Andean successional dry forests. This variability indicates that environmental heterogeneity does not manifest as a single pattern, but rather as a set of ecological responses adjusted to local conditions and land-use history. Water availability associated with precipitation was identified as the main variable structuring vegetation, accompanied by edaphic and thermal factors that vary between watersheds. Thus, environmental heterogeneity emerges as the dominant ecological mechanism that appears to regulate floristic composition.

The results partially confirm the hypothesis, showing that landscape transformation modulates, but does not by itself determine, floristic variation. Among the limitations of the study, it is recognized that the successional stages analyzed correspond mainly to early and intermediate stages. This is because, although some plots classified as late stages show structural features (such as greater basal area and height) typical of mature forests, their floristic composition still reflects early phases of succession. Therefore, detecting clear patterns of floristic differentiation likely requires long-term monitoring (≥ 20 years) that captures the full dynamics of tropical dry forest recovery. In this regard, tropical dry forest conservation and restoration strategies must integrate local edaphic and climatic conditions, prioritizing species with high drought tolerance that are adapted to their environmental context.

